# A Human-Machine Coupled System for Efficient Sleep Spindle Detection by Iterative Revision

**DOI:** 10.1101/454488

**Authors:** Dasheng Bi

## Abstract

Sleep spindles are characteristic events in EEG signals during non-REM sleep, and are known to be important biological markers. Manually labeling spindles by visual inspection, however, has proved to be a tedious task. Automatic detection algorithms generalize weakly for versatile spindle forms, and machine-learning methods require large datasets to train, which are unfeasible to acquire particularly for experimental animal groups. Here, a novel, integrated system based on a process of iterative “Selection-Revision” (iSR) is introduced to aid in the efficient detection of spindles. By coupling low-threshold automatic detection of spindle events based on selected parameters with manual “Revision,” the human task is effectively simplified from searching across signal traces to binary verification. Convergence was observed between resulting spindle sets through iSR, largely independent of their initial labeling, demonstrating the robustness of the method. Although possible breakdown of the revised spindle sets could be seen after multiple rounds of Revision, due to overfitting of the revised set to the initial human labeling, this could be compensated for by a Selection scheme tolerant to higher False-Negative rates of the machine labeling relative to the standard set. It was also found that iSR is generalizable to different datasets, and that initial human labeling could be substituted by low-threshold machine detection. Overall, this human-machine coupled approach allows for fast labeling to obtain consistent spindle sets, which can also be used to train machine-learning models in the future. The principle of iSR may also be applied for many different data types to assist with other pattern detection tasks.

**Significance Statement:** Electroencephalography (EEG) recordings are widely adopted in brain research. Abnormalities in the occurrence of particular EEG waveforms, such as sleep spindles, can be used to diagnose psychiatric diseases. Traditionally, human experts have labeled EEG traces for sleep spindles, a time consuming process; automated detection algorithms, however, often yield inaccurate results. This study introduces a new method for efficient sleep spindle detection with a human-machine coupled system that can iteratively revise labeled datasets, enabling convergence towards a robust, accurate spindle labeling. This system eases large-scale sleep spindle detection, which can yield datasets for both biological analyses and for training machine-learning models. Furthermore, the underlying method of iterative revision can be used to analyze other types of patterns efficiently.

## Introduction

Sleep spindles are 0.5-3s bursts in electroencephalography (EEG) recordings with central frequencies of 8-16 Hz and a distinctive waxing-waning pattern generated by the thalamic reticular nucleus (TRN) (Huupponen et al., 2000, 2007; Duman, 2005; Sitnikova et al., 2009). As a unique characteristic of non-rapid eye movement (non-REM) sleep in mammals, sleep spindles have been used as important biological markers in sleep research and for investigating the functional role of the TRN in memory consolidation and synaptic plasticity (Diekelmann and Born, 2010; Fogel and Smith, 2011). Furthermore, abnormalities in the density of sleep spindles has been experimentally determined to be correlated with schizophrenia, autism, and ADHD (Ferrarelli et al., 2007; Wamsley et al., 2012; Wells et al., 2016; Antony et al., 2018), among other psychiatric disorders. Thus, counting spindles in EEG recordings and determining their characteristics could have valuable applications in medical diagnostics.

Traditionally, sleep spindles have been manually marked by human experts (Warby et al., 2014; Purcell et al., 2017). However, this task is time-consuming and is difficult for large-scale studies. In light of this, several automatic detection algorithms have been developed, primarily based on signal processing techniques such as band-pass filtering, amplitude thresholds, or a variety of transformations (Schimicek et al., 1994; Duman et al., 2009; Devuyst et al., 2011; Adamczyk et al., 2015). However, the fine-tuning of algorithm parameters against human gold standards can be a laborious and unsystematic task. Furthermore, besides the difficulty to obtain large gold standard spindle sets, labeling by human experts may not be completely reliable, as reflected by large variabilities between manually labeled sets (Warby et al., 2014).

More recently, machine-learning methods have also been researched in to improve the performance of automated detection algorithms (Gorur, 2002; Ventouras et al., 2005; Camilleri et al., 2014; Ventouras et al., 2014; Tan et al., 2015). However, the training process is convoluted and often requires very large sets of human expert labels, which may be unfeasible to obtain for specific groups of subjects such as genetically modified mice. Moreover, the overfitting of machine-learning models may also be a concern when applying such trained models to new subjects.

This study presents a new method to address these issues by introducing an iterative approach for spindle detection, integrating both human labeling and algorithmic automatic detection in a process of “Selection-Revision” that systematically adjusts algorithm parameters. Starting with a short segment of manually labeled spindles, the algorithm processes the EEG data to obtain more potential spindle events, creating a larger label set which is then reviewed by human visual inspection. The revised label set is then used to perform parameter adjustment of the algorithm for better alignment, and the machine-detection-human-inspection process can be performed iteratively. For new datasets, the Revision process can start with generalized, low-false negative machine detection, eliminating the need for initial manual labeling. This system effectively reduces the human workload from a searching task to binary classification of spindles, and, aside from improving human consistency in labeling, can facilitate generation of large labeled datasets that can be used in future training of machine-learning models.

## Materials and Methods

### Obtaining of EEG recordings

The EEG data analyzed in this study was provided by Dr. Soonwook Choi at the Broad Institute of MIT and Harvard.

### Automatic spindle detection algorithm

An automatic spindle detection algorithm was implemented using MATLAB (R2018a, Mathworks, Inc.) based on the Short-Time Fourier Transform (STFT) (Gorur, 2002), achieving fast machine labeling of spindles (∼1 minute of running time for 6 hours of EEG recording). The EEG data was first preprocessed by smoothing and noise reduction. Previously sleep-scored non-REM segments of sleep were then transformed by STFT with a 300ms Hamming window and 250ms overlap. The power of the spindle frequency band (8-16 Hz) was calculated relative to the total power of the signal, and a double-threshold was applied on this power ratio for spindle detection. Segments 0.5-3 seconds long that crossed a lower threshold during their entire lengths and crossed an upper threshold at least once in their durations were considered spindles.

### Integrated interface

A custom, integrated interface was also developed using MATLAB for EEG data and sleep spindle visualization, manual labeling, and revision. In the interface, EEG data were displayed in customizable and scrollable lengths per screen, with the time axis labels marking 1-second increments and the vertical axis ranging from the minimum to maximum voltages recorded in the EEG data. Both preprocessed, non-REM segments and their corresponding filtered (band pass, 8-18 Hz) signals were displayed with linked time axes for reference.

For spindle labeling, human labelers could click on the start and end times of the spindle on the graph, and record the corresponding spindle events. Labelers were able to reference previously labeled events throughout one labeling session, and could modify their labels by deleting events or updating start/end points. Labelers could also continue with a new session by loading previously saved matrices containing the corresponding timestamps for labeled spindles. Initial labeling of a segment usually took 3-6 times the length of the EEG recording.

For fast spindle revision, reviewers could choose to accept, reject, or modify the start/end points for each potential spindle event shown, based on surrounding graphs of the EEG signal and the corresponding band pass filtered signal. Reviewers were blinded from the origin of the events shown (from the machine set, the human set, or both). Revision of one spindle usually took 2-3 seconds, and the time spent during revision depended largely on the size of the revision set.

### Performance evaluation

Since an overwhelming majority of the EEG recording does not correspond to any spindle events, a by-sample performance analysis would yield an extremely large TN, causing inflation of the overall performance measurements. Thus, for the purposes of this study, a by-event performance evaluation was adopted, categorizing each spindle event marked by either side being evaluated as one of the following: True Positive (TP), False Positive (FP), and False Negative (FN). These criteria, especially the FN rate, was used extensively when measuring the performance of various algorithm parameter pairs.

As not all marked spindles perfectly overlapped with each other, it was necessary to determine whether to implement a threshold for overlapping. However, it was found that, when comparing the machine labeled sets to the initial human labeled sets, nearly all spindle events had sufficiently large overlapping percentages. Moreover, this overlap was found to have increased in the following rounds of revision. Therefore, it was unnecessary to implement an overlapping threshold but rather to accept an event as TP as long as there existed an overlap of sorts between the two sets.

When comparing two manually labeled or revised sets that did not solely consist of machine-generated labels, it was meaningless to define spindle events as FP or FN. Thus, recall and precision between the two sets were calculated, where

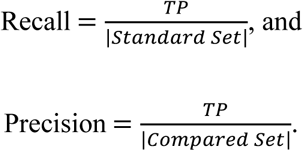

The harmonic means of recall and precision were calculated so that the F1 score would provide a measurement of the similarity of any two sets, and would remain independent of the order of the two sets.

### Statistical tests

Statistical analyses, in the form of t-tests (one-tailed, two-tailed, or one-sample test for mean), were performed using MATLAB. The significance thresholds used were *α* = 0.05 for (*), *α* = 0.01 for (**), and *α* = 0.001 for (***). Averages are plotted as mean ± standard deviation.

See **Table 1** for a summary of the statistical analyses used.

**Table 1.**
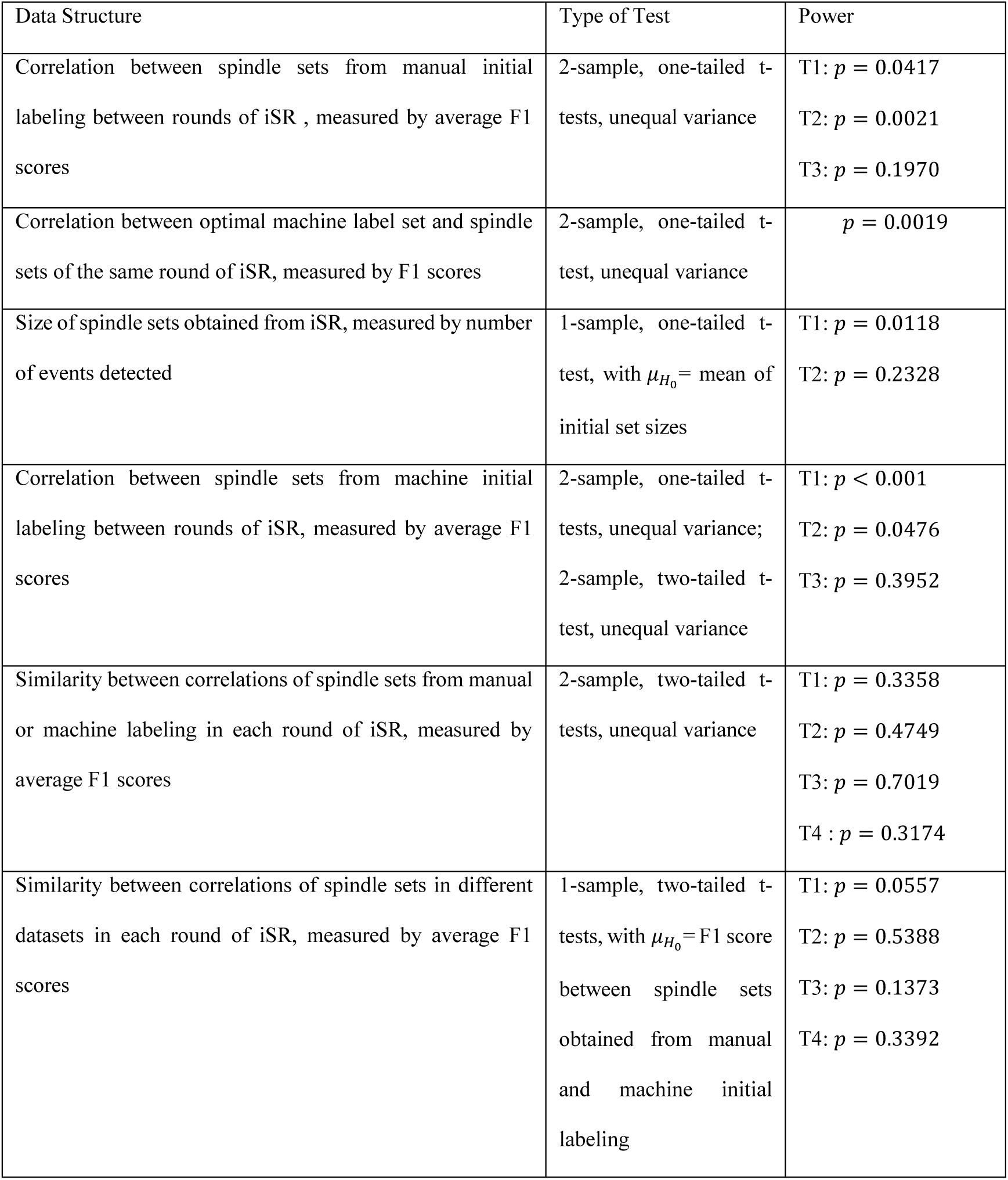
Summary of Statistical Analyses. Table shows a summary of all statistical analyses performed in this study. Columns denoting the type of data being analyzed, the type of statistical test used in the analysis, and the resulting *P*-values are shown.

| Data Structure | Type of Test | Power |
| --- | --- | --- |
| Correlation between spindle sets from manual initial labeling between rounds of iSR , measured by average F1 scores | 2-sample, one-tailed t-tests, unequal variance | T1: $p = 0.0417$<br>T2: $p = 0.0021$<br>T3: $p = 0.1970$ |
| Correlation between optimal machine label set and spindle sets of the same round of iSR, measured by F1 scores | 2-sample, one-tailed t-test, unequal variance | $p = 0.0019$ |
| Size of spindle sets obtained from iSR, measured by number of events detected | 1-sample, one-tailed t-test, with $\mu_{H_0}$ = mean of initial set sizes | T1: $p = 0.0118$<br>T2: $p = 0.2328$ |
| Correlation between spindle sets from machine initial labeling between rounds of iSR, measured by average F1 scores | 2-sample, one-tailed t-tests, unequal variance;<br>2-sample, two-tailed t-test, unequal variance | T1: $p < 0.001$<br>T2: $p = 0.0476$<br>T3: $p = 0.3952$ |
| Similarity between correlations of spindle sets from manual or machine labeling in each round of iSR, measured by average F1 scores | 2-sample, two-tailed t-tests, unequal variance | T1: $p = 0.3358$<br>T2: $p = 0.4749$<br>T3: $p = 0.7019$<br>T4 : $p = 0.3174$ |
| Similarity between correlations of spindle sets in different datasets in each round of iSR, measured by average F1 scores | 1-sample, two-tailed t-tests, with $\mu_{H_0}$ = F1 score between spindle sets obtained from manual and machine initial labeling | T1: $p = 0.0557$<br>T2: $p = 0.5388$<br>T3: $p = 0.1373$<br>T4: $p = 0.3392$ |

### Code Accessibility

The custom code described in the paper for automatic spindle detection, performance analysis, labeling, and Selection-Revision is freely available online at https://github.com/dashengbi/siris. For this study, the code was run on a Windows 10 computer with an Intel i5 CPU and 8.00 GB RAM.

## Results

### Integrated system for EEG analysis

The EEG data obtained was passed through a systematic process of manual and machine labeling, performance evaluation, and Revision, to repeatedly add to or delete from a “standard” set of spindles (See **Figure 1**). The iterative system for spindle detection is based upon two processes: Selection and Revision. Selection is the process during which machine sets with certain parameters (that can achieve appropriate alignment of algorithm-labeled spindles with those in the standard set) are chosen, and Revision is the process during which a large spindle set (including spindles from both humans and the machine) is reviewed and its spindles accepted or rejected. Selection can be applied either following an initial generation of a manual labeled set, or following a Revision process, and this Revision-Selection sequence may be applied iteratively to adjust both algorithm parameters and the standard set in order to achieve better consistency of detection.

**Figure 1.**
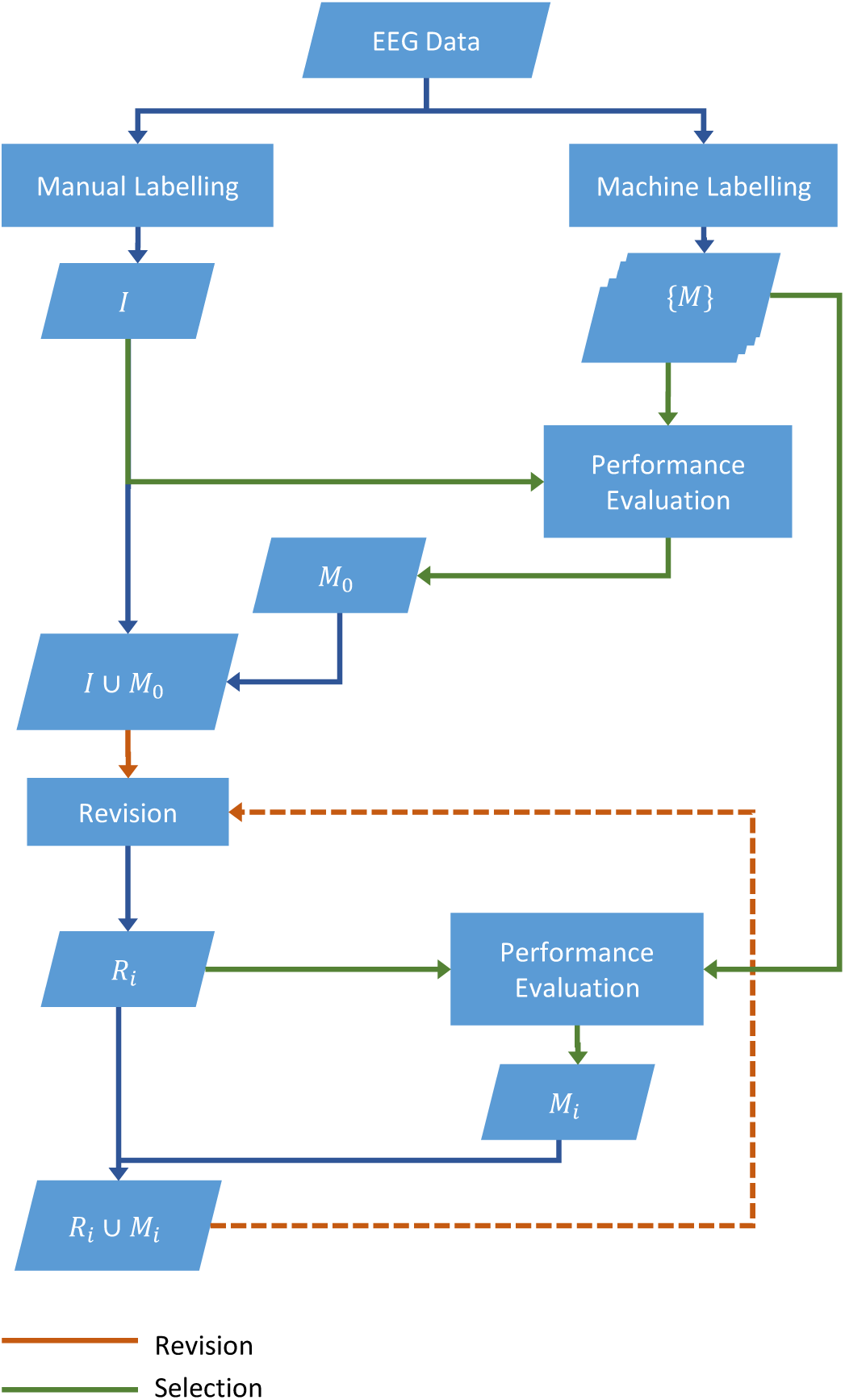
Diagram for iterative EEG analysis system. *I* is the spindle label set generated from initial human labeling (only needed once), {***M***} is the set of all machine label sets with different parameters, *M*_*i*_ is the machine set with parameters obtained from Selection (*i* ∈ *N* ∪ {0}), and *R*_*i*_ is the revised standard set after *i* rounds of Revision (*i* ∈ *N*^+^). During each round of Revision, spindles are either accepted or rejected; thus *R*_*i*+1_ is necessarily a subset of *R*_*i*_ ∪ *M*_*i*_.

### Alignment between different human and algorithm labeled sets

Upon obtaining several standard sets and machine labeled sets, the spindle events were cross-analyzed. Some sample spindle events labeled by different humans and by the automatic detector are shown in **Figure 2(a)**. It was found that although some characteristic spindles were labeled by most or all of the humans and the machine, the agreement between different detectors varied largely for other events. To evaluate the performance of the algorithm, machine-labeled sets with different parameters were compared with the standard sets, and plots specifying the recall versus precision rates of the machine labeled sets relative to one standard set were drawn (See **Figure 2(b)**). The outermost curve on the Recall-Precision plots represents the intrinsic tradeoff between the two statistics that the machine algorithm embodies. The plots of the agreement between different human sets are also shown. Initially, the agreement rates between different initial sets were lower than those with the machine set with optimal performance (shown by the point on the outermost curve furthest from the origin). After multiple rounds of revision coupled with their respectively selected machine sets, the agreement between revised sets from different initial sets increased greatly to points higher than those of the optimal algorithm sets.

**Figure 2.**
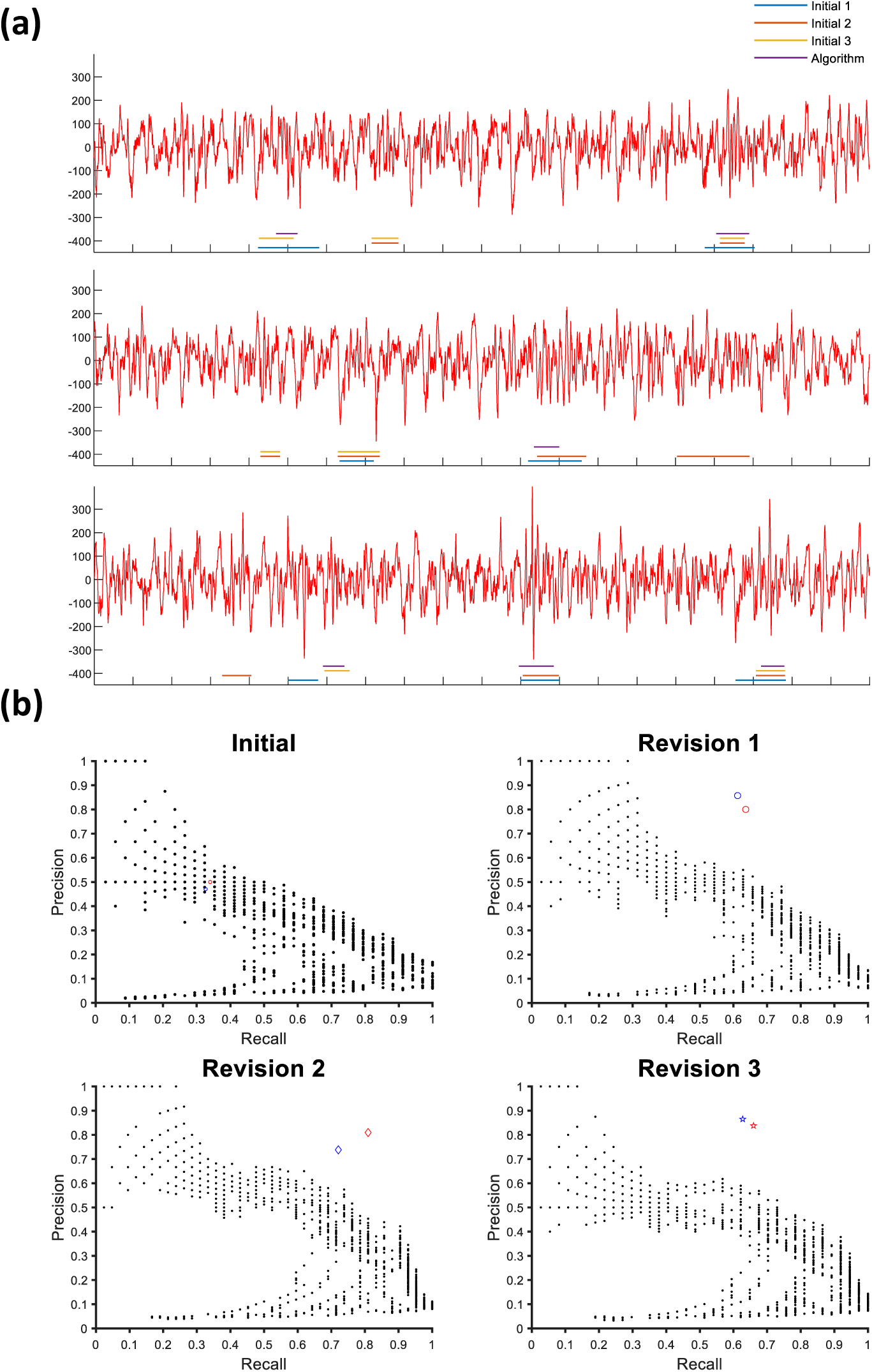
Recall-Precision Plots for Different Algorithm Parameters and Human Label Sets. **(a)** Several spindles in three different initial human labeled sets and an algorithm set optimized for performance on one of the human sets are shown. **(b)** Each point on the plot corresponds to the recall and precision of one spindle set relative to the standard set from one human labeler after different rounds of revision. Black points are results from machine labels with different parameters, and red/blue points refer to revised spindle sets starting with different initial spindle sets.

### Selection of machine labeled set for Revision

To select an appropriate machine set for Revision, both false negative (FN) and false positive (FP) rates needed to be considered. The respective FP rates and machine label set sizes relative to various FN thresholds (so that the FN rate of the system would not exceed such a threshold) were plotted. Higher FN rates would cause the iterative Revision system to have more inherent false negatives relative to the ground truth, as the algorithm may not be able to detect certain spindle events the human marked as negative. Higher FP rates were correlated with much larger sets of spindles for humans to review, thus decreasing the efficiency of the Revision system. During revision, if the machine set with the optimal F1 score was selected, on average, the first round of Revision would have a FN rate of 0.5069—that can also be seen as approximately the rate of spindles not detected by humans that would also be neglected by the machine (See **Figure 3(a)**). Previous studies have shown that the FN rates between different human experts are around 0.25-0.3 (Warby et al., 2014); thus it would suffice to use a machine labeled set with FN rate <0.2 to cover most spindles in the ground truth. The mean number of spindles in a 1200s segment of non-REM sleep detected using the optimal-F1 score machine parameters was 53, while the mean number of spindles detected by machine algorithms while limiting their FN rate to less than 0.2 was 147 (See **Figure 3(b)**).

**Figure 3.**
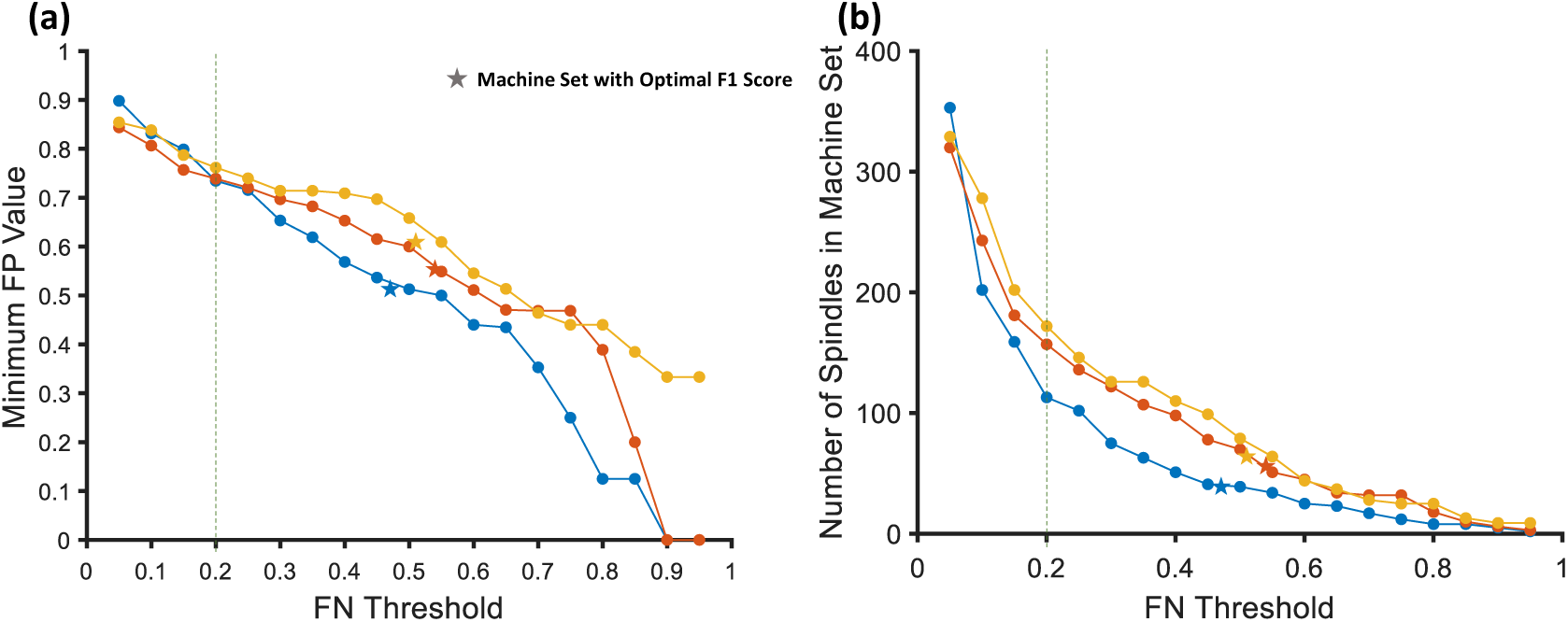
Tradeoff Between False Positive Rate, Time Efficiency, and FN Rate. Three different initial non-REM segments of 1200s length, labeled by different human experts, are shown in different colors. The stars represent the statistics obtained for the respective machine sets with the optimal F1 score performance. **(a**) Minimum FP value achieved by algorithm while maintaining FN rate below certain thresholds. **(b)** The number of spindles in machine labeled set for each of the corresponding FN thresholds. The number of machine-labeled spindles is directly related with the time required for Revision.

### Significant increase in correlation between differently obtained spindle sets upon Revision

It was found that iterative Revision could increase the alignment between resulting spindle sets from largely different initial human labeled sets (See **Figure 4(a)**). A measurement of the alignment between different spindle sets at a given time could be obtained by calculating the average F1 score between all possible pairs of spindle sets. A measurement of the disparity between all spindle sets could be obtained by calculating the standard deviation of the F1 score between all possible pairs of spindle sets. Using a one-tailed t-test assuming unequal variances between different revision rounds, it was found that the average cross-compared F1 score significantly increased as a result of the first round of Revision (*P* = 0.0417). The average cross-compared F1 score significantly increased between the two subsequent rounds of Revision (*P* = 0.0021). However, the average cross-compared F1 scores were not significantly different between the second and third rounds of Revision (*P* = 0.1970). Thus, by applying iterative Revision, the agreement between spindle sets labeled by different humans could converge to one standard set.

**Figure 4.**
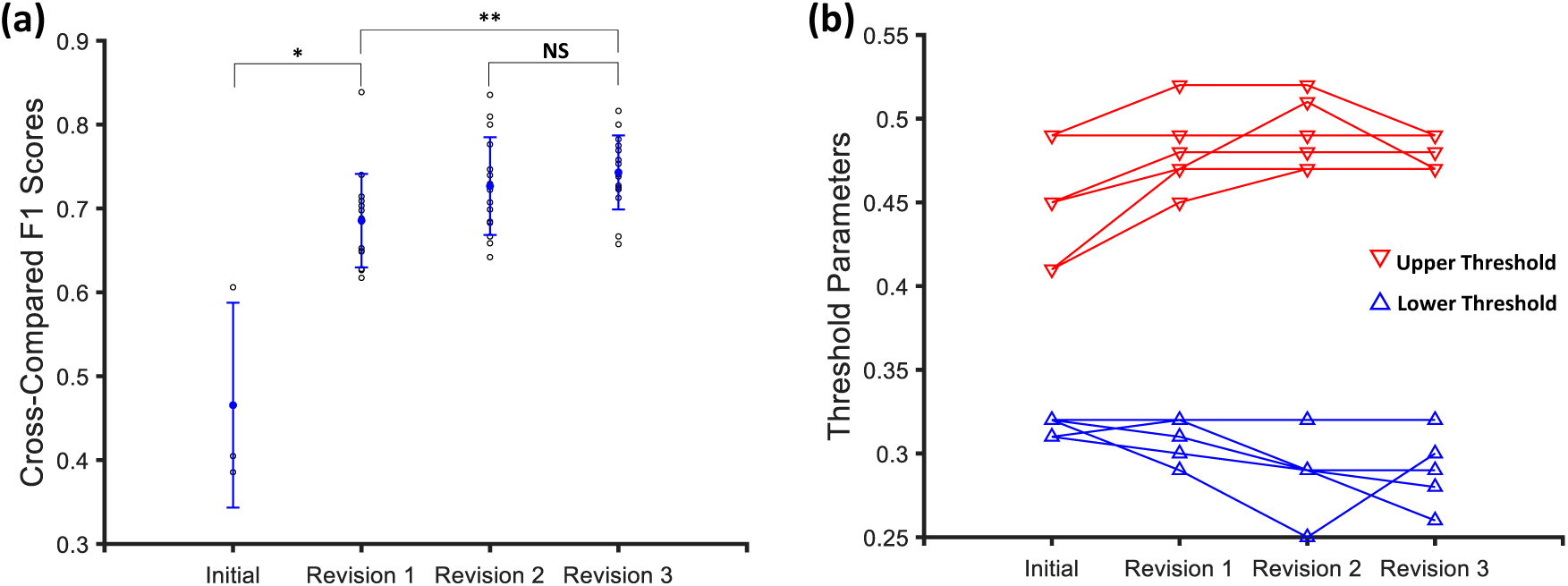
Convergence of Cross-Compared F1 Scores and Upper Threshold After Multiple Rounds of Revision. **(a**) Scattered circles show the F1 score between all possible pairs of initial sets or revised standard sets under two different Selection schemes (FN < 0.2 or Optimal F1 score). There were 3 points in the initial round and 21 points in all following rounds of Revision. Blue filled points show the average cross-compared F1 scores in a given round, and blue lines show the standard deviation of cross-compared F1 scores in that round. **(b)** Scattered triangles show the values of the “threshold” parameters of the optimal algorithm parameter sets selected with respect to the standard sets after various times of Revision.

As a result of the increased alignment between spindle sets, the algorithm parameter sets selected using iteratively revised sets also showed signs of convergence (See **Figure 4(b)**). In particular, the upper threshold of the algorithm showed a decreasing trend in standard deviation as Revision was applied continuously.

### Overfitting of F1 score and breakdown of standard set with high-FN Selection schemes

For two Selection schemes, three rounds of Revision-Selection were performed, starting with three different initial sets. The F1 scores obtained for the Revision rounds were found to be significantly different (See **Figure 5(a)**), with those for the Selection scheme of choosing the optimal machine set (as measured by F1 score relative to the standard set obtained from each previous round) being higher (one-tailed t-test, *P* = 0.0019). Examining the size of the revised standard sets obtained after several rounds of revision (See **Figure 5(b)**), it was found that there was a significant decrease in the revised set sizes relative to the sizes of the initial human-labeled sets (one-tailed t-test, *P* = 0.0118). For the standard sets obtained by Revision after Selection with a FN threshold of < 0.2, there was not a significant deterioration of the set size (one-tailed t-test, *P* = 0.2328). Therefore, Selection schemes with higher FN tend to introduce overfitting of the standard set with machine sets, and can cause revised standard sets to significantly decrease in size, deviating away from the ground truth; it is necessary to adopt a Selection scheme with low FN to utilize the Revision process fully.

**Figure 5.**
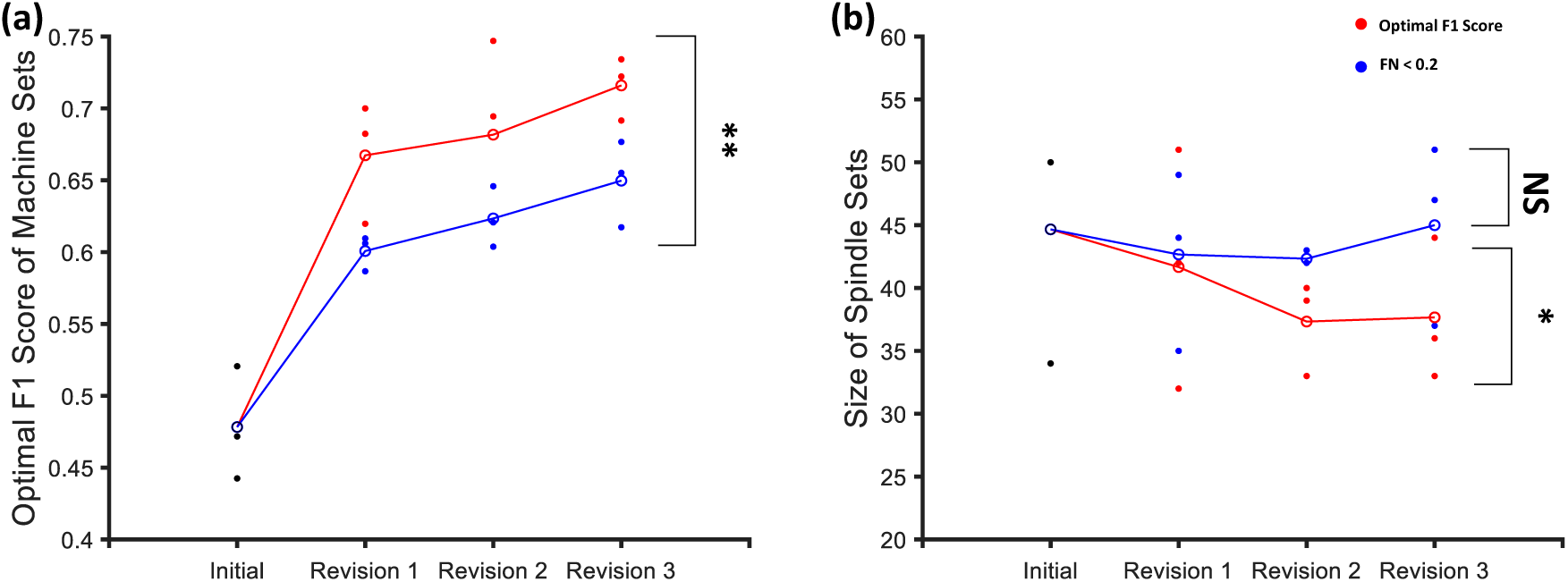
Optimal F1 Score of Algorithm-Labeled Sets After Multiple Rounds of Revision. **(a**) Points show the optimal F1 scores that the algorithm can obtain through parameter adjustment (with respect to differently obtained initial human sets or revised sets during the same round of Revision). The connected circles show the trends of the average performance of the algorithm within each Selection scheme changing over time. **(b)** Points show the sizes of the initial or revised standard sets (*I* or *R*_*i*_, where *i* ∈ {1, 2, 3}). Connected circles show the trends of the average sizes of the revised standard sets under different Selection schemes.

Indeed, by iteratively applying Revision with Selection scheme of *FN* < 0.2, it was found that the agreement rate between revised standard sets from different initial sets steadily increased (See **Figure 6**).

**Figure 6.**
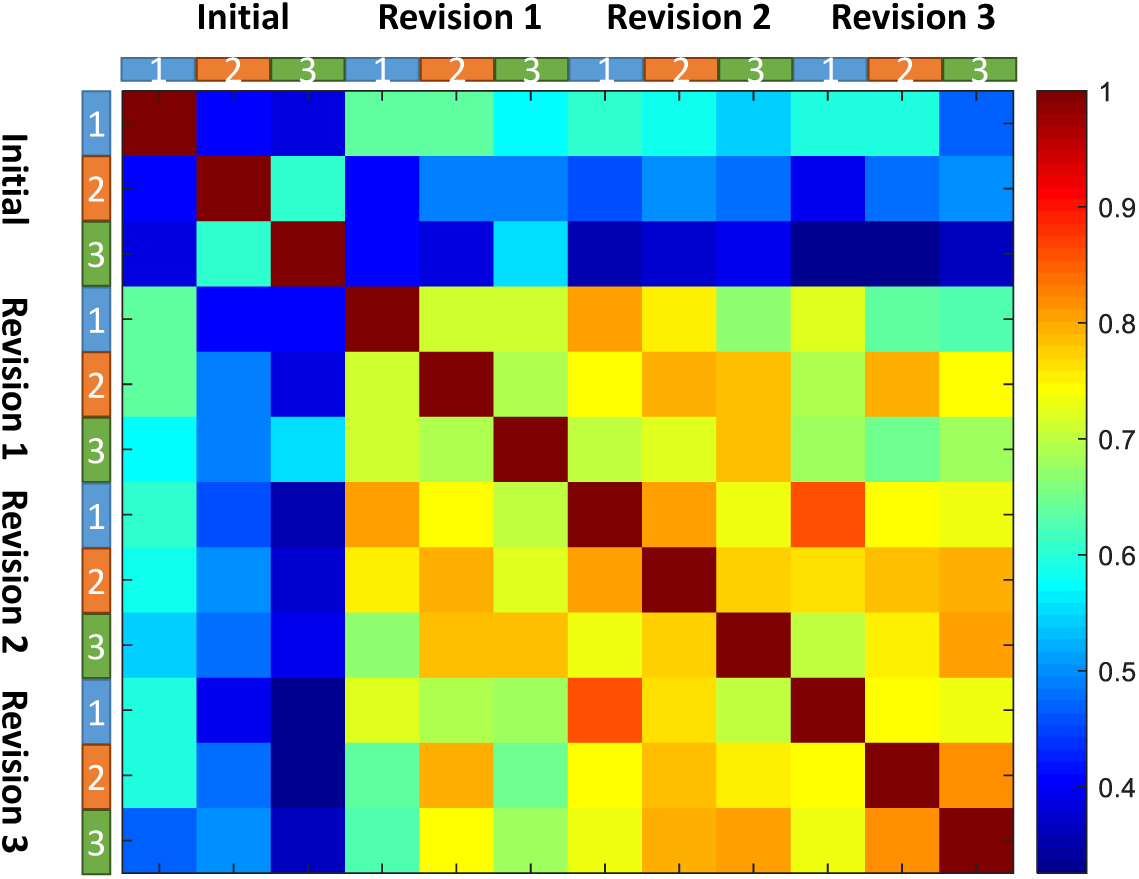
Increased Agreement between Standard Sets with Different Initial Sets. Heatmap with color coding showing three different sets in their respective rounds of Revision. Each scaled grid shows the agreement (F1 score) between two spindle sets.

### Revision is generalizable to extended datasets by applying algorithms with Selected parameters for initial labeling

To test whether pure algorithm labeling could be applied to novel datasets with minimal loss of spindles, separate rounds of Revision, both on the previously mentioned dataset and a new dataset, were performed with the initial labeling being generated using the automatic detection algorithm. The parameters of the algorithm (lower threshold = 0.21, upper threshold = 0.47) were determined based on the previously obtained standard sets with manual initial labeling after three rounds of Revision so that the FN rate of the initial algorithm set would not exceed 0.1. The correlation between the revised sets from initial algorithm and manual labeling were obtained (See **Figure 7(a)**).

**Figure 7.**
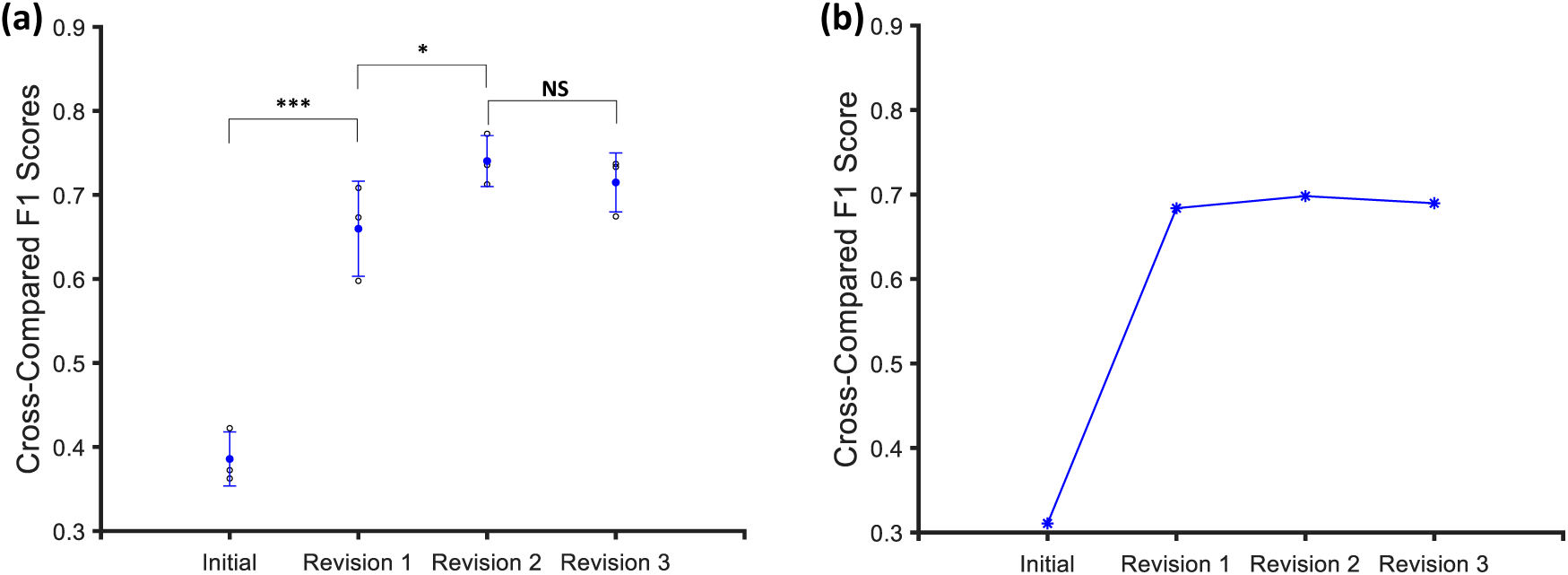
Convergence of Revised Spindle Sets Starting with Algorithm Labeling. **(a)** Points show the F1 score between a standard set revised with initial human labeling and a standard set revised with initial algorithm labeling. Blue filled points and lines show the average F1 score of the algorithm initial/revised set relative to human initial/revised sets of the same Revision round. **(b)** Points show the F1 scores between a standard set revised with initial human labeling and a standard set revised with initial algorithm labeling for a different trace, with the algorithm parameters used being the same as those in (a).

It was found that Revision greatly increased the alignment between standard sets revised from initial algorithm and manual labeled sets (one-tailed t-tests: *P* < 0.001 from Initial to Revision 1 and *P* = 0.0476 from Revision 1 to Revision 2). After two rounds of Revision, the agreement rate did not change significantly (two-tailed t-test, *P* = 0.3952). Furthermore, it was found that for each round of Revision, the mean agreements between the revised sets obtained from initial algorithm labeling were not significantly different from those obtained from initial human labeling (two-tailed t-tests: *P* = 0.3358, *P* = 0.4749, *P* = 0.7019, and *P* = 0.3174 for the initial round, and the first three rounds of Revision, respectively).

To further demonstrate the generalizability of the convergent nature of Revision to other datasets of EEG, three rounds of Revision (with Selection scheme of FN<0.2) were performed on another dataset using two different initial sets (manual labeling and machine labeling). By comparing the F1 scores between the two (See **Figure 7(b)**), it could be seen that the agreement rates increased through repeated Revision. Moreover, it was found that for each of these rounds of Revision, there was no significant difference in the F1 scores between the standard sets from two methods of initial labeling across the datasets (two-tailed t-tests: *P* = 0.0557, *P* = 0.5388, *P* = 0.1373, and *P* = 0.3392 for the four rounds, respectively). Therefore, the machine labeled initial sets, when combined with iterative Revision, are able to generate reliable standard sets. Thus, the method can generalize well to novel datasets.

## Discussion

It was found that after a process of iterative Selection-Revision to adjust initial human labeled spindle sets with the introduction of certain machine-detected events, the agreement rates between different standard sets improved greatly. It was also found that a FN rate threshold was necessary for effective adjustment of the initial human set, as higher FN would cause deterioration of the spindle set size after several rounds of Revision.

The system of iterative Selection-Revision improves the quality of standard sets resulting from an initial set that need not be very carefully labeled by human experts. Though generally, the FP and FN rates of machine detection were high, once an appropriate machine set (with a controlled FN rate) was combined with a manually labeled set, spindle events that were previously undetected by humans could be noticed during revision. Furthermore, the Revision process also limits the introduction of false-positives into the system, as each spindle event is subjected to multiple rounds of scrutiny.

The most pressing issue that the Selection-Revision system addresses is that of time. Previously, accurate labeling of sleep spindles required laborious searching of EEG traces by human experts (Ventouras et al., 2005); with the system of Selection-Revision, an initial set without such time-consuming manual labeling may be applied and revised iteratively until the resulting standard set evolves towards the ground truth. During Revision, human validation of machine-labeled spindles is much easier to perform as compared to manual detection. For a 20-minute long segment of non-REM EEG, applying two rounds of Revision requires only around 10 minutes, while estimated times for careful manual labeling spindles in such data can be as long as 2 hours (based on the author’s own experience). This indicates that iterative Revision may reduce the human workload by as much as tenfold.

However, there are several aspects that should be taken into consideration. These include the potential bias of the human reviewer, who may be inclined to label and revise spindles with inconsistent standards, and the inherent FN/FP rate tradeoff of the Revision system, caused by the limitations of the machine detection algorithm (Ventouras et al., 2005; Tan et al., 2015). These concerns may be addressed by introducing certain “confusion” spindle events to evaluate the possibility of human bias during Revision, and by implementing more algorithms that are able to analyze different aspects of the EEG signal. By combining these algorithms that focus on distinct spindle characteristics, it would be able to provide more accurate machine-augmentations, reducing the human workload even more. Ultimately, the human-machine coupled Selection-Revision system may generate large training sets at a fast speed, thus facilitating machine-learning models for large-scale spindle detection.

Despite convergence being observed between revised spindle sets and between selected upper threshold parameters of the algorithm, convergence of the lower threshold was not observed. This may be caused by the complex relationship between algorithm parameters and spindle sets, and that automatic detection may be more sensitive to upper thresholds than to lower thresholds. More systematic tests are needed to determine whether such a relationship exists. For practical purposes, however, two rounds of Revision is often sufficient to obtain reliable gold-standard spindle sets.

In conclusion, this study has introduced a novel method for efficient sleep spindle detection based on a mechanism of iterative machine-augmented human Revision. It has shown that through multiple rounds of Revision, largely different spindle sets that were initially coarsely labeled by humans could evolve and converge into standard sets more closely aligned with each other. This method of iterative Selection-Revision can be applied as a systematic means of fine-tuning automatic detection algorithm parameters with the absence of a meticulously generated gold standard, and can also be used to expedite the process of gold-standard label generation for training machine-learning models of spindle detection.

## Acknowledgements

This study was started during a summer internship at the McGovern Institute for Brain Research at MIT. The author thanks Dr. Guoping Feng for providing the internship opportunity and scientific guidance; Dr. Zhanyan Fu for patient supervision; Dr. Soonwook Choi for teaching me about EEG and sleep spindles, and for providing EEG recordings of mice from the Broad Institute of MIT and Harvard; Xuyun Sun for helpful discussions on automatic detection algorithms; and Tina Naik for performing sleep-scoring of EEG traces. Initial sleep spindle sets used in the study were extracted from manual labels performed together with Dr. Soonwook Choi and Xuyun Sun. The author would also like to thank his parents for their support, encouragement, and suggestions; and all the members of the Feng Lab at MIT for a stimulating discussion environment.

